# Disease relevance and replicability of deep learning gene expression prediction

**DOI:** 10.64898/2026.09.24.753534

**Authors:** Ada Zhang, Shinya Tasaki, David Connell, Bernard Ng, Chris Gaiteri

**Affiliations:** Department of Psychiatry, SUNY Upstate Medical University, Syracuse, NY, USA; Rush Alzheimer’s Disease Center, Rush University Medical Center, Chicago, IL, USA; Department of Neurology, Columbia University Irving Medical Center, New York, NY, USA

**Keywords:** sequence-to-expression models, deep learning, genomic architecture, reproducible science

## Abstract

Recent deep learning (DL) models predict average gene expression levels from DNA sequences with high overall correlation to measured values. We examine these DL models through the lens of disease research. The seemingly high overall performance of DL models is largely due to capturing whether genes are “On” or “Off”, and to a lesser extent, disease-relevant expression level changes for “On” genes. Indeed, the more bimodal the gene expression distribution, the better the reported performance. We track the extent of this issue across tissues, molecular systems, and cancers. Compounding the model evaluation issue, we find inconsistencies between the published code and reported performance, highlighting the importance of versioning and publishing performance evaluation code. These findings indicate that the high reported performance of popular DL models falls unexpectedly short in disease applications, and that the problem of personalized genomic prediction remains far from solved in a disease context.

## 1 Introduction

The majority of genetic risk for most common complex diseases accumulates from variants with small effect sizes that are widely distributed across the genome [1–6]. The disease variants are overwhelmingly non-coding and often affect the expression levels of nearby genes (expression quantitative trait loci – eQTL’s) [7, 8]. These variants contribute, alongside environmental effects, to moderate changes in mRNA expression in common complex diseases: typically in the 10–50% range versus healthy controls [9–12], with higher multiples in some cancers [13–15]. This diffuse disease structure motivates the desire to know how every DNA nucleotide could increase or decrease gene expression. Such an account of non-coding variants would be the beginning of a causal narrative of disease, which would help explain how subsequent regulatory effects, such as transcription factors [16] and chromatin state [17], generate disease effects [18–21]. Indeed, drugs targeting molecules with genomic disease variants are more likely to be approved [22, 23]. However, providing this genome-wide understanding of the actions of millions of variants, in a realistic human cellular environment, far exceeds current experimental capacity.

In contrast to the extensive experimentation previously required to investigate the function of a single variant, sequence-to-expression deep learning (DL) models try to build a universal “grammar” to interpret any known or novel variant. At its theoretical limit, such a grammar could predict effects for any basepair in any person. As increased input context for LLM’s generally improves performance [24, 25]; reported performance for these methods has improved from the original 1000-basepair context [17] to the present thousand-fold higher 1-megabase input [26], and as they include additional epigenomic data types [26–28]. Recent DL approaches have reported variance explained (R^2^) of ∼0.8 and higher, which are approaching the ceiling of technical replication, and would seem to indicate the expression prediction problem is largely solved [27]. If these methods are modeling the effects of the non-coding variants so accurately, how can there be any question about their utility for disease research?

Interpreting or predicting the effect of person-specific variants helps drive interest in these models, but it was never their sole purpose. However, their broader performance has generated interest in this question, with some recent studies indicating that sequence-to-expression models offer scant or no predictive power for personalized disease research [29–31]. Recently developed methods take this lack of performance on personalized genomes as a given, and directly attempt to improve it by fine-turning the base sequence-to-expression models [32–35]. Those results indicate a continuing barrier from across-gene prediction, vs. utilizing those models for person-specific variant interpretation, although there are signs that increasing the number of individuals in such studies may yet improve that aspect [34]. In light of the surprisingly variable performance appraisal of sequence-to-expression models, we conduct additional investigations to help determine their potential for disease genomics applications.

To resolve these divergent assessments, we consider the core logic of machine learning in genomics: that improved model performance results from better representing real biological processes [36]. This logic depends on two assumptions: that better performance metrics are driven by meaningful aspects of biological data, and that this performance itself is replicable. We find that these assumptions do not hold for major sequence-to-expression deep learning methods. The performance metrics for gene expression from Enformer often do not relate to meaningful aspects of disease biology, wherein expression changes are moderate. The method mainly predicts whether a gene is “On” or “Off”, but not precise gradations within expressed genes, as would be ideal for complex disease, as a function of genetic variation. We explore the extent of this limitation across multiple tissues, in multiple molecular domains, and with simulations, which indicate this limitation is broadly applicable to subsequent methods utilizing the same widely used datasets. Considering the second assumption, that performance must be replicable, Enformer does match published performance, albeit with these limitations on relevance. However, another popular method, Basenji, in our hands, does not match published performance when utilizing released code. Also, running their code with data augmentation enabled did not reach reported performance on human data. In light of these significant limitations, we propose biological and software engineering guidelines for all future deep learning genomics methods, to more robustly utilize their potential.

## 2 Results

### 2.1 Bimodal expression distributions influence performance metrics

High correlation between sequence-predicted gene expression and actual mRNA levels contributes to interest in deep learning genomics models. Enformer is able to predict gene expression with correlations that are “approaching experimental-level accuracy, estimated at 0.94” [27]. A closer inspection, however, will indicate that a bimodal distribution of “On” versus “Off” gene expression is a major contributor to this high correlation. Such binary classification bears little similarity to how mRNAs are regulated in disease, wherein significant case/control differences are often entirely composed of genes that are “On” in both conditions [37–39]. Nevertheless, a major intended application of this method is “improved diagnostic tools for diseases of genetic origin.”

To first demonstrate the bimodal distribution, we use the test partition of Basenji’s released data and examine the predicted versus ground truth gene expression for the Basenji [28] and Enformer [27] models (Fig. 1). Gene expression values were extracted from raw data following the protocol described in [27] (see methods). Both models show overall Pearson correlations of *R >* 0.8. The underlying distribution is strongly bimodal, with a peak of “Off” genes centered around -1.5 and a peak of “On” genes centered around 1 (normalized units). (“Off” and “On” genes had an average read count of 1.2 and 141.8, respectively.) If we separate the genes into “Off” and “On” genes according to their ground truth values (dotted line in Fig. 1) and compute the correlation of predicted expression to ground truth expression within each group, we see that performance drops substantially for both Basenji (*R*_OFF_ = 0.53, *R*_ON_ = 0.53) and Enformer (*R*_OFF_ = 0.53, *R*_ON_ = 0.64).

**Fig. 1:**
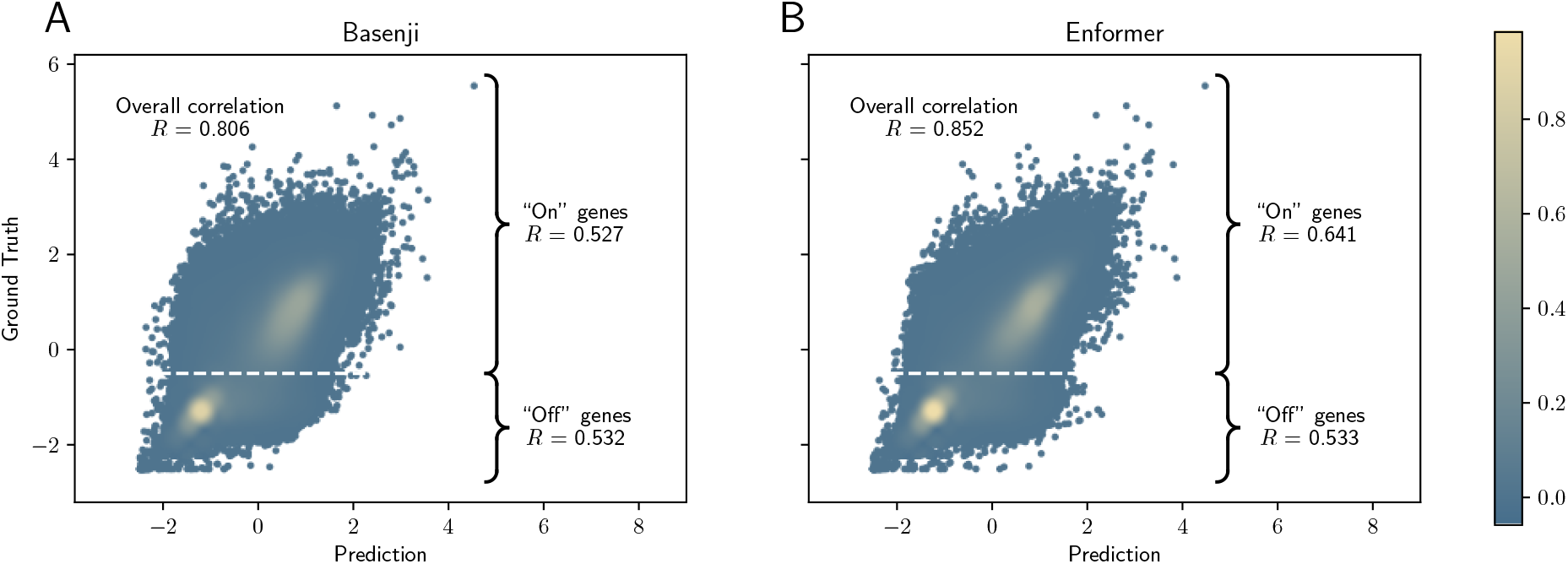
Predicted vs. ground truth gene expression for (A) Basenji and (B) Enformer. Each point represents the expression of one gene in one CAGE track. Ground truth values are obtained from the test partition of Basenji’s released data [28] and then processed using Enformer’s protocol [27] to obtain ground truth gene expression values. The same protocol is applied to model outputs to obtain predicted gene expression levels. Points are colored according to density, with blue representing low density, and yellow representing high density.

It may be possible to provide more meaningful performance metrics by examining the “On” and “Off” gene populations separately. To do this robustly, for each cap analysis of gene expression (CAGE, see methods) track, we vary the threshold value for separating the groups and compute the correlation of the genes with expression *>* threshold (“On” genes, Fig. 2B) and those with expression *≤* threshold (“Off” genes, Fig. 2C). We then average the correlation across the 638 CAGE tracks at each threshold value. The corresponding average percentage of samples (genes) across CAGE tracks with ground truth expression values above and below a given threshold value are shown in Figs. 2D and E respectively. Because Basenji and Enformer share the same ground truth data, their values align exactly for these two subpanels. By cross-referencing with Enformer’s distribution (Fig. 2A), we show that correlations within the “On” and “Off” gene groups do not approach the overall correlation until the threshold value starts to include samples from the other mode (Fig. 2D). Fortunately, Enformer’s improvement over Basenji stems largely from improved correlation within the “On” genes.

**Fig. 2:**
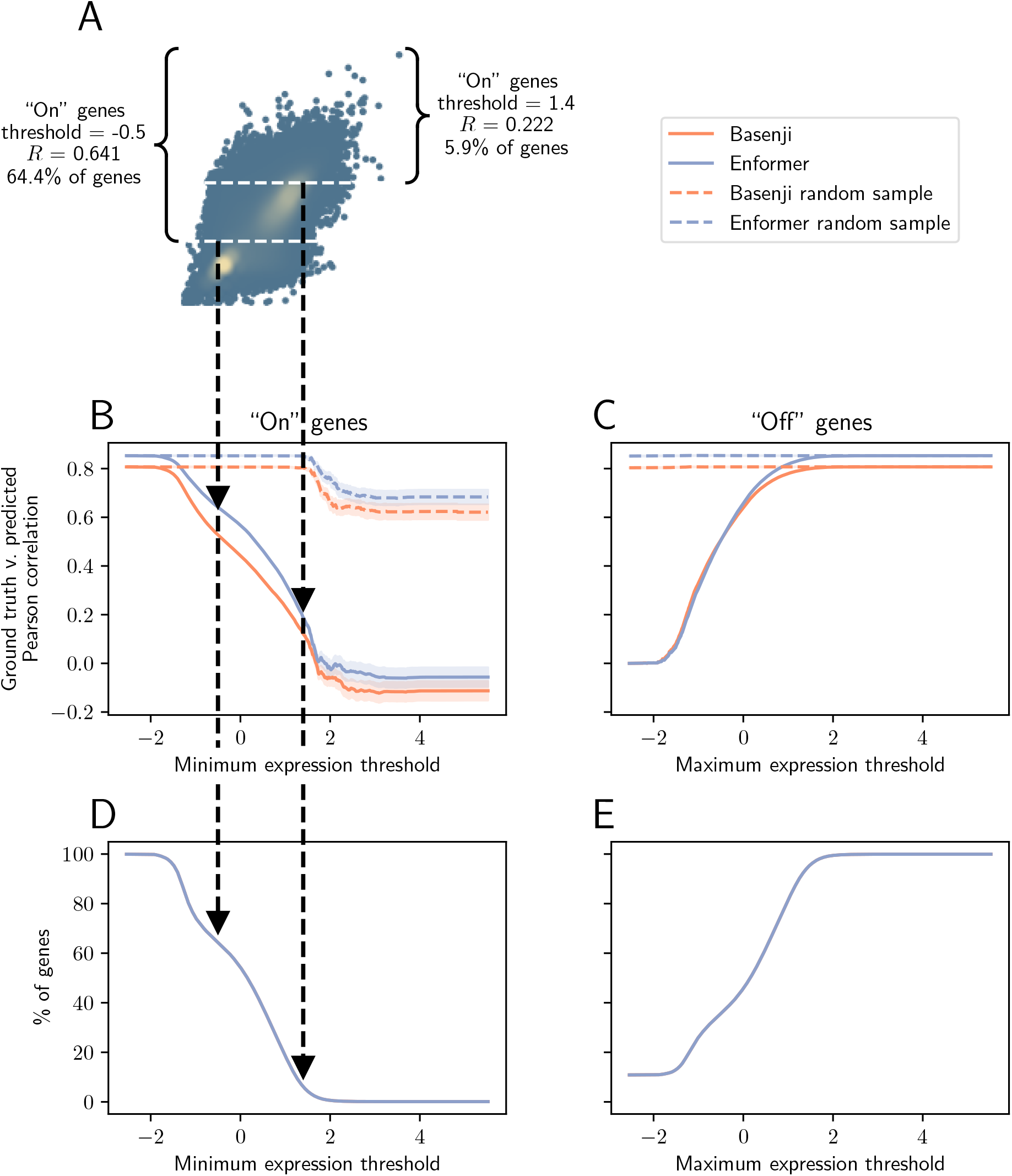
Correlations of “On” and “Off” genes for varying threshold values. A) Enformer predicted and ground truth gene expression (as shown in Fig. 1B) with example thresholds shown at two differing values (one at -0.5 and one at 1.4) which determine a given set of “On” genes and the corresponding Pearson’s correlation *R* and % of genes within that grouping. B) The dotted black lines link the threshold lines in panel A to their corresponding correlations and the % of genes above that threshold value (panel D). Shaded area indicates standard error. C) In a parallel analysis, we also consider all possible thresholds for defining “Off” genes. D) The percentage of genes defined as “On” decreases as the threshold rises. E) The percentage of genes defined as “Off” increases as the threshold rises.

To confirm that these findings were not due to the smaller sample size of “On” and “Off” genes compared to the overall set of genes, we repeated this analysis with a random sampling of all genes (Fig. 2B-C, dotted lines). In particular, for a given threshold, we compute the number of genes *N* within each group, randomly choose *N* samples from all genes, and then compute the correlation across these *N* genes. The dotted lines reach the overall correlation value once *N >* 100 (corresponding to *∼* 5%), indicating that the reduced correlations we observe in the “On” and “Off” gene groups are not due to small sample size.

To further investigate the effect of bimodality on correlation, we used a Gaussian Mixture Model (GMM) to identify the “On” and “Off” groups of ground truth expression within each CAGE track. We then measured the distance between these two peaks and compared it to the Pearson correlation of ground truth versus predicted expression within that track (Fig. 3A bottom row). CAGE tracks with more distinctly bimodal distributions (i.e., larger distance between GMM modes) were associated with higher prediction correlation (Basenji *R* = 0.577, p *<* 1e-57, Enformer *R* = 0.567, p *<* 1e-54). For robustness, we also remove the 1% most extreme values (points marked by in *×* Fig. 3A) and the association remains (revised Basenji *R* = 0.553, p *<* 1e-49, revised Enformer *R* = 0.540, p *<* 1e-47). The point of these results is not to compare Enformer and Basenji, but rather to show that both methods are strongly susceptible to this effect where greater bimodality is associated with higher model performance, across genes and CAGE tracks.

**Fig. 3:**
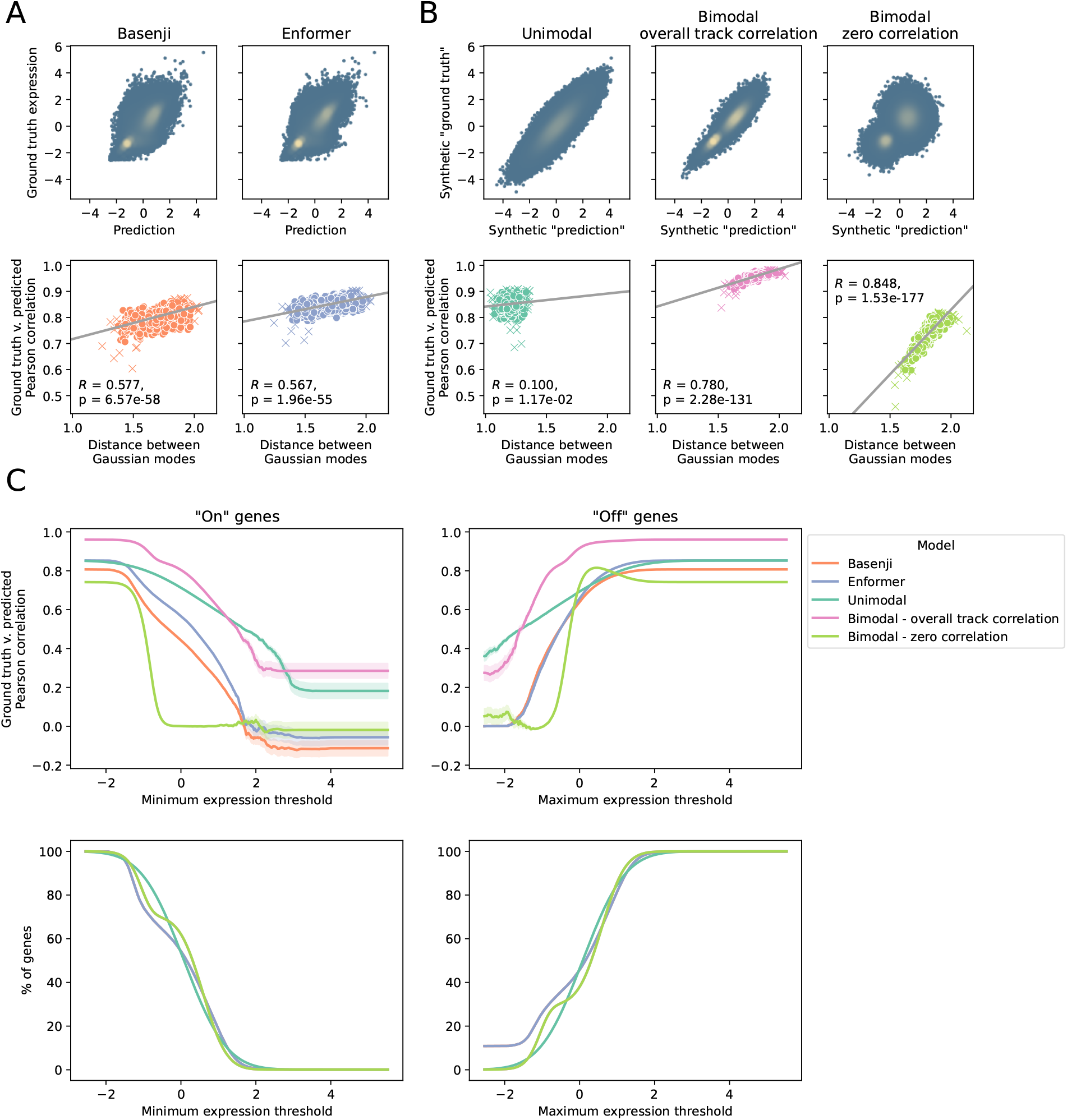
Extent of bimodality correlated with apparent veracity of predictions. A) Top row: Density plots showing the distributions of Basenji and Enformer. Points represent one gene in one track. Bottom row: Using distance between means of “On” and “Off” gene groups of a CAGE track (estimated via a Gaussian Mixture Model) versus Pearson correlation between ground truth and predicted expression from Basenji. Points represent CAGE tracks. B) Parallel density plots (top row) and “On” / “Off” gene group distance versus Pearson correlation (bottom row) for the three synthetic datasets. Left: Unimodal distribution matches the overall mean and correlation of Enformer data to ground truth expression. Middle: Bimodal distribution matches the means of the modes of Enformer and the overall track correlation is applied to each mode. Right: Bimodal distribution still matches the means of the modes of Enformer, but each mode has zero correlation with ground truth. See methods for details. C) Correlations of “On” and “Off” genes for varying threshold values. Top left: Analogous to Fig. 2, we define “On” genes according to various thresholds and test how that set correlates with ground truth, for various real and synthetic models. This test is performed for the complete possible range of cutoffs (x-axis). Top right: In a parallel manner, we consider the correlations to ground truth for just the “Off” genes. Enformer’s results were used to compute the track correlations, which is why the Unimodal correlation (teal line) merges with that of Enformer (blue line) once all samples are included within the group. Bottom left: In the process of testing every possible threshold, for increasingly stringent thresholds, the number of “On” genes decreases. Bottom right: In the converse of the bottom left, as the threshold rises, the number of “Off” genes increases.

We generated three synthetic datasets based on Enformer performance to better understand how the correlations of different distributions may contribute to performance metrics (Fig. 3B top row). The “unimodal” model considers a type of null comparison case in which we match the overall mean and correlation of the distribution shown in Fig. 1B, but without the two modes prominent in the real data. This enables us, in further testing, to understand how much bimodality may be contributing to apparent performance. We also explored two bimodal distributions: one modeled after the correlations between predicted and actual expression within the “On” and “Off” genes, and another with no correlation within those modes (see methods). This latter bimodal distribution can be seen as a null model of performance that would be expected with no predictive power within the “On” genes, beyond that they are highly expressed.

As with Basenji and Enformer, we used a GMM to identify the “On” and “Off” groups of genes within each synthetically generated track. We then compared the distance between these groups (as a measure of bimodality) to the overall correlation of that track (Fig. 3B bottom row). The “unimodal” distribution (Fig. 3B left) shows almost zero correlation between GMM distance and Pearson correlation, which represents an ideal scenario where model performance is independent of the bimodality of the underlying distribution. Meanwhile, the “bimodal zero correlation” distribution (Fig. 3B right) demonstrates a strong trend between bimodality and correlation, supporting the hypothesis that it is easier to achieve high correlation on bimodal datasets.

We then compared these synthetic datasets to the Basenji and Enformer results by applying the same performance test of varying the threshold and computing the correlation within each group, as described above. These results are shown in Fig. 3C. As before, “On” gene correlations are shown in Fig. 3C top left, “Off” gene correlations in Fig. 3C top right, and the corresponding average number of samples for a given cutoff value are shown in Fig. 3C bottom row. Recall that ground truth values are used to compute the % of genes for a given threshold, which makes Basenji and Enformer align exactly in Fig. 3C bottom row, but synthetic “ground truth” expression values are randomly generated, leading to slight differences in their plots. Notably, the bimodal distribution using zero correlation for each mode (green line) still achieves an average correlation across tracks of 0.742 when all points are included, which demonstrates how bimodality can lead to misleadingly high overall correlation even if a model is only predicting “On” and “Off.” The bimodal distribution using the overall track correlation for each mode (magenta line) achieves even higher correlation when both modes are included, further supporting our finding of the effect of bimodality on correlation.

Finally, we compared performance of these continuous models of gene expression to two types of “Naive” (binary) models that only predict “On” or “Off” expression. These binary models are not useful from a disease perspective because they cannot predict moderate changes in expression. However, comparing their performance to their continuous counterparts offers insight into the weaknesses of current performance metrics. Fig. S3 illustrates how we generated the two Naive models using Enformer’s data as an example: One had access to ground truth expression data, predicting “On” if ground truth expression was greater than a threshold of -0.5, and “Off” otherwise (Fig. S3A), and the distribution of this model’s predictions is given in Fig. S3B. (Note that, for the synthetic datasets, “ground truth” was the synthetically generated y-axis data.) This model represents the best possible performance by a Naive predictor and is represented by square markers in Fig. 4. The other “Naive” prediction model used a threshold of -0.5 on the predicted (Basenji, Enformer, or synthetically generated) expression values to determine “On” or “Off” (Fig. S3C-D). This model represents a lower bound of performance if a model were trained to only predict “On” or “Off” as opposed to continuous expression values and is represented by triangle markers in Fig. 4. These Naive models are compared to their continuous counterparts, represented by stars, in Fig. 4.

**Fig. 4:**
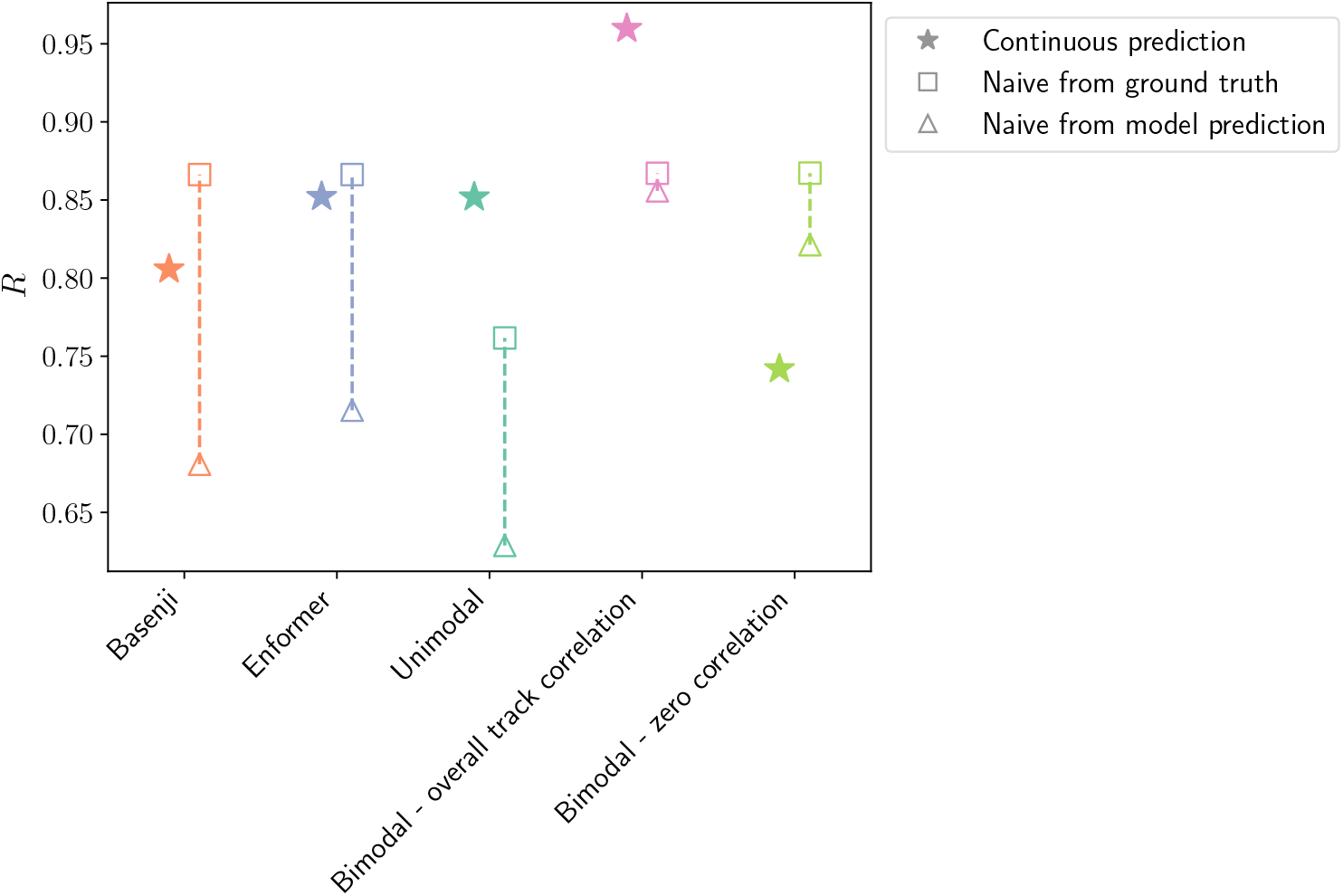
Comparing *R* of the continuous predicted value of the “models” versus a naive predictor, which only predicts “On” or “Off.” We expect the continuous models (stars) to have better *R* over the binary naive predictors (squares and triangles). The “Unimodal” and “Bimodal - overall track correlation” synthetic distributions demonstrate this expected behavior. Meanwhile, Basenji and Enformer, despite outputting continuous values, have worse performance than the Naive model built on ground truth data (square). Additionally, the “Bimodal - zero correlation” distribution demonstrates the degenerate case where the continuous model (star) is worse than its binarized version (triangle).

The general expectation is that a binary (naive) model’s performance will be strictly worse than that of a continuous model, particularly for disease applications where subtle variations in gene expression are key. Fig. 4 shows this expected behavior from the “Unimodal” and “Bimodal - overall track correlation” distributions. However, we see that Basenji and Enformer have worse performance than the Naive model built on ground truth data. While such a perfect Naive model is unrealistic to obtain from training, we remark that this ideal “On”/”Off” predictor is simultaneously useless for disease understanding and yet has better Pearson *R* than Baseenji and Enformer, highlighting the weakness of Pearson *R* as a metric for gene expression prediction performance. Additionally, the “Bimodal - zero correlation” distribution demonstrates the degenerate case where the continuous model is worse than its binarized version.

### 2.2 Disease-focused performance analysis

With a similar strategy as above, we examined the effects of bimodality on prediction for each tissue. Each CAGE track represents a sample obtained from a particular tissue. We manually separated the CAGE tracks into groups with the same, or highly similar, tissues of origin, so that we could perform tissue-level analyses. Non-tissue-specific samples were excluded from analysis, as were tissues with fewer than 3 samples. Some samples were drawn from cancerous tissue, so we report results both including and excluding cancer cells from tissue groups. Fig. S1 shows tissue-based model correlations and GMM distances. ANOVA tests demonstrated significant differences in correlation and GMM distance between tissues, whether we included or excluded cancerous samples (Table S1). We also computed GMM distance within a tissue to test its effect on correlation. We used two strategies to generate tissue-based GMM distances: (1) we averaged the GMM distances and Pearson correlation for all CAGE tracks from a given tissue, see Fig. S2A, and (2) we combined the ground truth values of all CAGE tracks for a given tissue and fit a new GMM to that distribution and computed a new Pearson correlation between the combined ground truth and predicted values, see Fig. S2B. In both cases, there is a significant positive correlation between bimodality (measured by GMM) and model performance (measured by Pearson correlation) when cancerous samples are included. When cancerous samples are excluded, the effect is lessened, possibly due to smaller sample size and some tissues being excluded due to all samples being cancerous.

We performed gene set enrichment analysis (GSEA) to better understand model performance across different tissue sets. We split the CAGE tracks into three groupings: (1) cancerous and non-cancerous, (2) different tissues (43 groups), (3) different tissues separated by cancerous and non-cancerous (72 groups because not all tissues had cancerous and non-cancerous samples). For each grouping, the median absolute error of Basenji and Enformer were calculated for each group. We found a slight positive correlation between gene expression and error, therefore we normalized the error by performing a linear fit of the ground truth expression to error and saving the residuals. GSEA was then performed on this normalized error (Tables S2-S17). We examined the GO categories with significantly elevated or degraded prediction accuracy.

GO categories related to olfactory receptor (OR) genes (n = 44, in this dataset) had negative NES, indicating the median absolute error was low, and the models had good performance for these GOs (Tables S9 and S15 for Basenji and Enformer respectively). As an example, GOBP “Detection of Chemical Stimulus” was significant in 72 of 72 tissue/cancer groups with an average p-value *<* 1e-5 and p-value *<* 1e-4 for Enformer and Basenji respectively. Biological details of olfactory receptor genes may be relevant to the high prediction accuracy for this family. Olfactory receptors directly respond to molecules in the environment, generating neuronal activity that ultimately drives olfactory perception [40] of a trillion odors [41] through unique activation patterns across hundreds of receptors [42]. These typically G-protein coupled receptors [43] are activated directly by airborne molecules, triggering cAMP-calcium intracellular cascades. However, every tissue assayed for ORs has shown expression of at least one OR and up to several hundreds [44] where they perform diverse sensing roles including: apoptosis [45], chemotaxis [46], secretion [47] cell-cycle [48], adhesion [49], inflammation [50] with multiple signaling cascades (PLC/IP3, MAP/ERK and RhoA). Despite the diverse functions across tissues, and that neurons typically only express a single OR gene per cell [43], in this data we observe that the average correlation between each pair of OR genes across all tracks has a mean *R* = 0.789 (permutation p *<* 0.001, Fig. S4). The gene set expression is overall lower than expected (permutation p *<* 0.001, Fig. S4). Possible explanations for the high accuracy on this group is the similarity of promoter sequences, which have *∼*20-60% sequence similarity [51, 52]. As expected from this, olfactory gene promoters tend to share some transcription factors—they are overall enriched for EBF1, LHX2 [53] and MEF [54] binding sites. In light of the moderate sequence similarity, low expression, and highly significant correlations in expression among all ORs in this data, the finding that OR gene expression is the most predictable should be interpreted with caution. The high prediction accuracy for this large family genes is likely not equivalent to a collection of independent predictions on genes with diverse sequences and diverse expression patterns.

Utilizing differential GSEA results for cancer vs non-cancer data sources, across all tissues, we find that GOs related to transcription factors were positively enriched among genes with poor predictions for both Basenji and Enformer (Tables S3 and S5 respectively). Many transcriptional categories were significantly differential—for instance, “GO MF transcription regulator activity”, p *<* 0.01. Origins of poor predictions for TFs in cancer necessarily stem from either alterations in their promoter sequences or altered expression dynamics. Enformer uses reference data so we do not have access to the whole genome sequence of each data type. While there are some indications or alterations in promoter sequences of TFs in cancer [55, 56], there is substantially more evidence for cancer effects on TF expression levels through altered regulatory changes via enhancers [57, 58], histone acetylation [59] and altered DNA methylation [60]. All of these epigenetic systems comprise part of the physical mechanisms through which promoter sequences have their effects, and ostensibly are embedded within the sequence-expression relationships the model learns in predominantly healthy-state data.

Indeed, Enformer can be used to predict some epigenetics and derivative methods that include relevant enhancer sequences alongside promoter can improve performance [61]. In a cancer state, promoter-enhancer interactions may undergo wide-spread rewiring, which could reduce the relevance of relationships driven by control-state data. In light of the lower predictability of some gene classes in cancer, likely due to genomic alterations, it might seem surprising that there are also gene categories with improved predictions in cancer. In particular, sets related to immune response (p *<* 0.01) are more predictable in cancer tissues (Tables S3 and S5). The likely origins of this are that cancer tissues are included in Basenji and Enformer training data, and some relationships might only exist in those data, and conversely are not seen in control data. Subsetting to the samples in which these relationships exist naturally generates higher predictability. The other broad biological class that is more accurately predicted in cancer samples (p *<* 0.01) are those related to olfactory receptors and related subprocesses, such as G-protein coupled receptors (Tables S3 and S5). Olfactory receptors were also the gene set that was overall more predictable across every tissue (p *<* 1e-6), so this result indicates they are particularly predictable in a cancer context, potentially due to a range of non-olfactory actions in multiple aspects of cancer [62]. Overall these observations suggest in cancer samples alterations in TF promoters have effects that are not predictable, while variants in the promoters of immune system and olfactory receptor genes have effects that are more predictable, at least with these methods and data sources.

We also examine if gene predictability has effects that go beyond any particular class of gene by testing if genes that behave predictably attract scientific interest. Therefore, we tested if predictability of gene expression was linked to the popularity of that gene in scientific literature. To do that, we calculated the frequency of a gene being mentioned in all of Pubmed, and if that had any relation to its predictability (see methods). We computed the Spearman correlation of log_10_ publication frequency and model performance. For Basenji, there was no significant correlation between model performance and publication frequency. For Enformer, however, we found a slight correlation (p = 0.003), indicating that the performance improvements from Enformer over Basenji pertains to genes that are of higher interest to the scientific community Fig. 5.

**Fig. 5:**
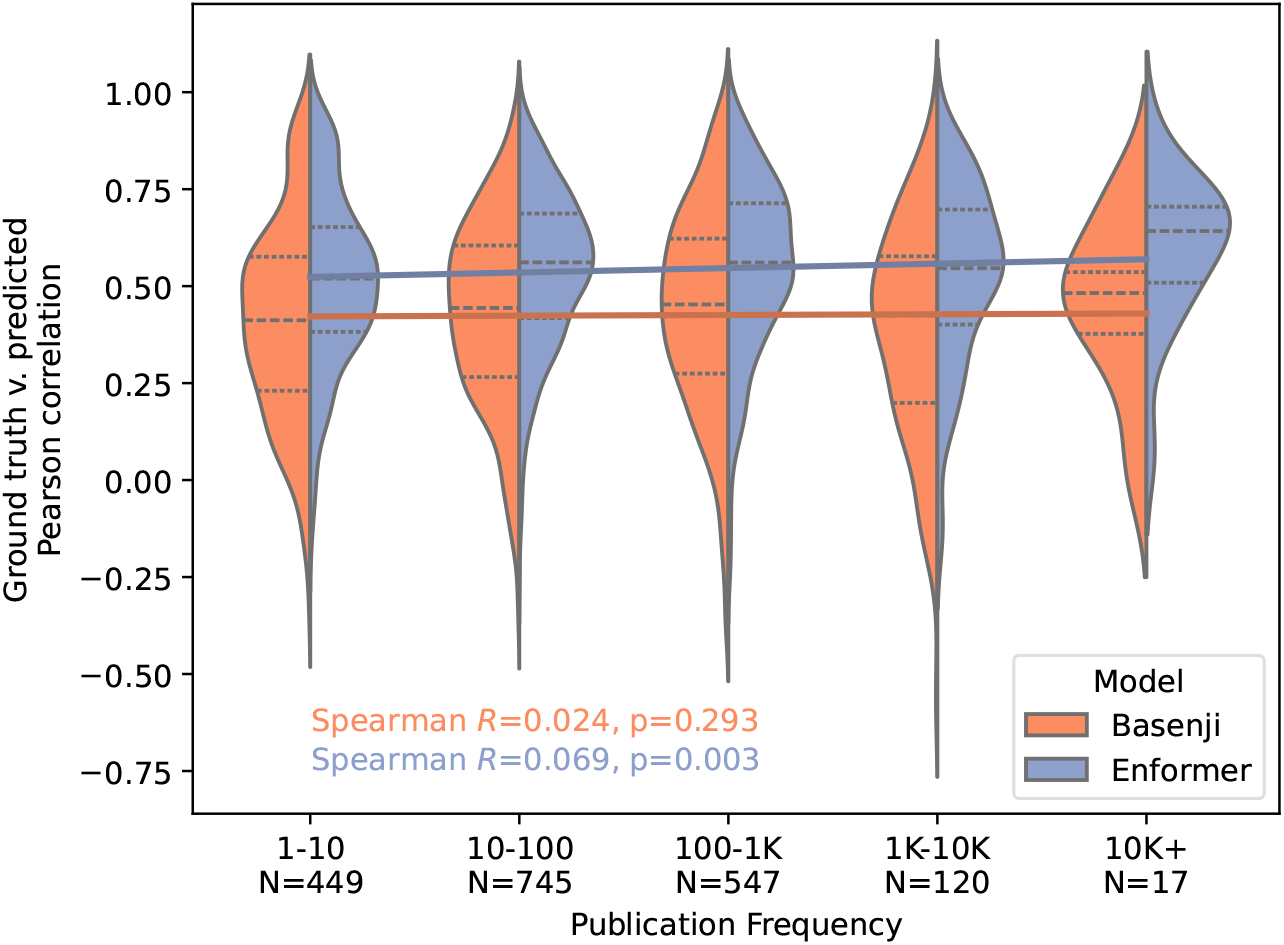
Gene publication frequency versus model performance. Model performance distributions (and the corresponding number of genes) are shown for each publication frequency bin.

### 2.3 Architecture performance is often not reproducible

While model performance might not be as practically useful as the high correlation values imply, it should at least match published results. To attempt to confirm this, we downloaded the pre-trained Basenji model and also tried to train a model from scratch using the code and dataset provided, and building our virtual environment using the prespecified `environment.yml` that was also released. We also ran inference on Enformer’s pre-trained model using their released code and environment `requirements.txt` as reference. Performance of Enformer’s pre-trained model did match what was reported in the paper, which incidentally also validates our performance calculations Fig. 6. However, the pre-trained model released by Basenji had lower performance (*R* = 0.626) than their reported “human & mouse trained” and “human-trained” performances (*R* = 0.645 and 0.632, respectively).

**Fig. 6:**
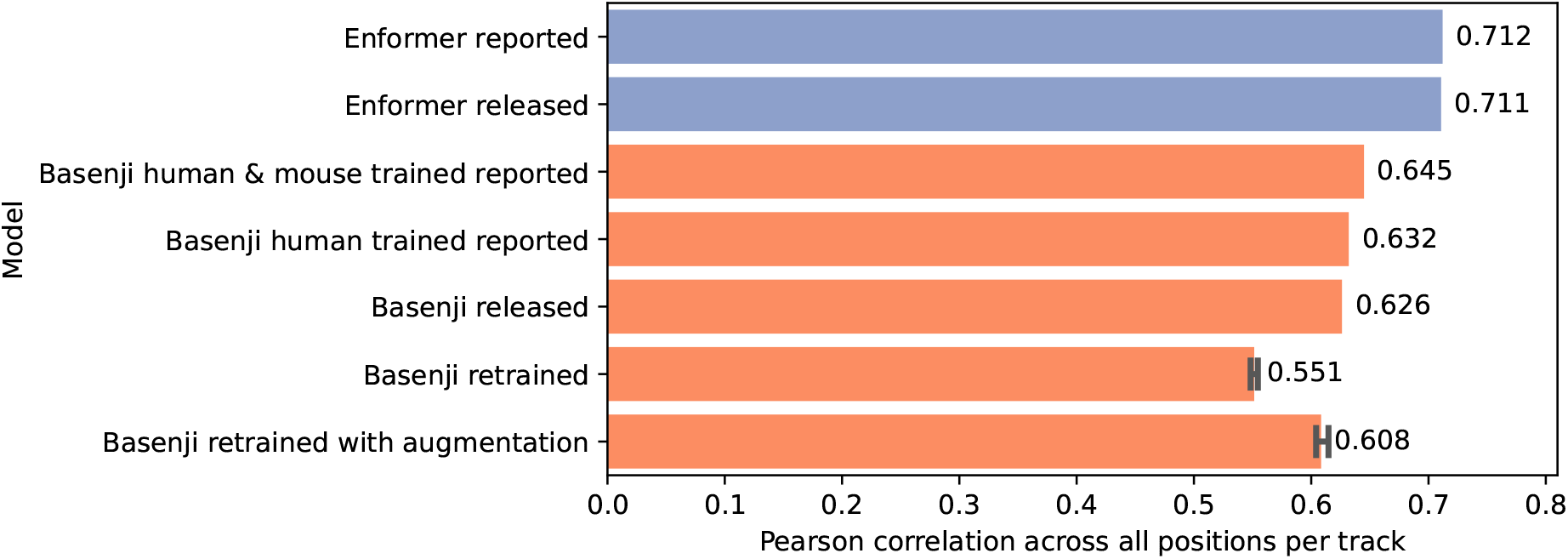
Overall correlation comparison between reported performance, released models and retrained models. Reported performance values are the correlation values reported in the publication. Released model performance was obtained by downloading the released, pretrained models, running inference, and then computing performance. Retrained model performance was obtained by downloading the code and data used to train the models, retraining from scratch, and then running inference and computing performance. This procedure was repeated four times to compute performance variance due to stochasticity in training. Inspection of the released parameter file indicated that it did not perform the data augmentation procedures described in the publication, so the parameter file was edited to perform data augmentation and the model was retrained four times again.

Training models from scratch leads to some variability in performance due to the stochastic nature of training, although we expect model performance to be close to reported performance values.

However, using Basenji’s provided code and data did not replicate their reported performance, reaching only *R* = 0.551 compared to the reported human-only trained performance of *R* = 0.632 (Fig. 6). Replicate experiments indicate that model training is quite stable, with the four runs having *R* within 0.547-0.555. Comparing the published code with the released parameter file, which specifies the architecture (number and size of layers, etc.) and hyperparameters (learning rate, momentum, etc.), indicates the released code does not implement data augmentation of *±*3bp shift and stochastic reverse complement, although the paper reported this augmentation procedure. After detailed inspection of their training code, we modified the released parameter file so that data augmentation would be performed during training, but the final performance still did not match the reported *R* = 0.632. Instead, the four runs had a mean *R* = 0.608 with range 0.603-0.618 (Fig. 6). While a difference of 0.024 might appear minor, the performance improvement in Basenji—claimed to be due to data augmentation from multiple species—was 0.013 in humans [28], indicating such performance increments can be considered meaningful.

A likely remaining source of the discrepancy is in the architecture or hyperparameters, as specified in the parameter file. In general, values in the parameter file match what was reported in the paper with the exception of patience: the paper reported waiting for 30 epochs of no improvement, whereas the parameter file indicates a patience of 16. Momentum and learning rate values (0.99 and 0.15 respectively) were not reported in the paper, so it is unclear whether there are additional errors in the parameter file that lead to reduced performance after training.

### 2.4 Post-processing requirements are an impediment to tracking model advancements

Enformer reported significantly improved correlation values (from 0.712 to 0.849) by focusing performance calculations around gene transcription start sites (TSS), rather than calculating correlations across the whole track. However, this performance assessment requires processing multiple data files and cross-referencing between them, making it non-trivial to compute. As a single example of the processing required before performance can be estimated, for Enformer, one needs to:

1. Download GENCODE gene annotations, process to extract TSS of protein coding genes.
2. Cross-reference the gene TSS from GENCODE with the Basenji `sequences.bed` file, which describes the start and end points of each Basenji sample.
3. Convert the gene TSS into an index in the Basenji sample (a vector of length 896), for which CAGE tracks are binned into 128-bp bins, and 64 bins (8192 bp) are cropped from either side of the original 131,072 bp sample.
4. Collect all bins containing any TSS associated with a given gene, and their neighbors to either side, removing duplicates for TSS of the same gene that are in adjacent bins, and then sum for both ground truth and predicted gene expression values. (Note: gene expression predictions are obtained by running inference 8 times on the same sequence augmented with *≤*3 bp shift and reverse-complementation, and then averaging the predictions.)
5. Perform a log_10_(1 + *x*) transformation (log base not specified in text) and then normalize each ground truth CAGE track to zero mean and unit variance. Apply the same normalization parameters of each CAGE track to the predicted values.
6. Compute correlations across genes within CAGE tracks and average across tracks.

Thus, in this case and in general, the performance assessment procedure is far more complex than computing a simple correlation between two vectors. Errors in any part of this implementation—failure to filter for only protein-coding genes, incorrect conversion from gene location to sample index, failure to perform a log_10_(1 + *x*) transformation, e.g.—can lead to substantially varying results. Furthermore, some aspects of this implementation were not clearly described, such as which gene annotation version was used, which base was used for the log transformation, and whether duplicate bins were removed when summing gene expression within *±* 1 bin around each TSS.

In summary, while this performance assessment is very insightful on current gene expression prediction capabilities, it is difficult to implement, making it difficult to reproduce the findings of [27] and compare future models along this performance metric. For example, Avsec et al. [27] mentioned an interesting finding: experimental accuracy of gene expression is estimated at 0.94 based on replicate CAGE experiments. This experimental accuracy may be less dependent on the bimodality of gene expression, as compared to the model predictive performance, but this hypothesis is difficult to interrogate because the code for testing replicate performance was not released. Although the implementation of these replicate accuracy tests is briefly described, re-implementing these experiments would be complex and likely to be incomplete without further description, requiring at least downloading raw FASTA files from FANTOM, preprocessing them following the method described by [28], splitting the replicates into the two pseudogroups according to the Supplementary Table 1 of [27], and only then computing correlations between the two groups across genes.

## 3 Discussion

Stratified distributions, such as the gene expression datasets used by DL models, can have unexpected effects on reported correlations, as we have observed. The difference in performance in predicting all genes vs just “On” genes can be seen as a variant of “Simpson’s paradox” [63]. Based on published results of Basenji and Enformer, as well as simulations, we find the real–predicted correlations are largely driven by two distinct modes, in a manner unrelated to most disease applications, which are concerned with smaller fluctuations of genes that are expressed in both health and disease. In the case of gene expression, differing from the classic case of Simpson’s, the direction of the correlation does not reverse, but is simply much lower among the “On” genes than expected from the overall correlation of “On” and “Off” genes. This leaves substantial variation in “On” genes unexplained, which limits the utility of these methods for understanding actions of non-coding variants in disease. It also supports the concept [64] that it may be necessary to explicitly focus on person-to-person variation of genes in disease, vs trying to derive a universal “grammar” across all genes, from which disease effects within genes naturally follow. Research in that vein can improve personal genomic predictions, but there is a general challenge in producing some method that generalizes to unseen individuals and genes [32–35].

The increasing power and popularity of deep learning approaches to long-standing questions in biology has also generated new technical hurdles to participation. It is increasingly expensive to reproduce results, both in terms of the hardware required, and also the labor needed for preprocessing data and ensuring that the computing environment and versioning are correct. Recently, emphasis on “dockerization”, in which program environments are encapsulated, can significantly reduce wasted time chasing down clashing software versions that arise even with a simple attempt to reproduce major results. There is a level of replication that goes beyond Docker, in which you hash all datasets and code bases, in which you can prove that a particular code version has produced a particular result. Packages are available to support this rigorous provenance [65, 66]. However, achieving perfectly reproducible results across hosts systems is a challenging goal that few attempt.

The current state of computational biology is far from such replicability—merely reaching published performance levels is not guaranteed, as we have observed in this specific case of DL genomics.

Hurdles to analyzing and using other’s work can be greatly reduced by following a minimal set of guidelines that acts as a golden mean between the effort of designing perfect reproducibility and the simplicity of supplying only a `requirements.txt.` This minimum spec entails that all parameter files are actually made available, that they match published values, and actually reproduce specific figures. While model architecture code is often released, performance assessment is no longer comprised of simple arithmetic. In light of challenges in both interpretation and reproducibility, we propose three guidelines for future deep genomics publications. Related suggestions have been made [67–69], but following through on them is important for research efficiency, and something for editors and reviewers to insist upon. Formulated as a checklist, experience with deep learning sequence-to-expression models suggest that in the future care should be taken to:

1. Perform tests on the appropriate variables. This means focusing on expressed genes when the intended application is complex disease, and not conflating between-gene performance to within-gene performance.
2. Reproduce key individual figures with published code. Focusing on figures and end-to-end reproducibility, rather than the mere existence of (possibly incomplete) parameter files, forces all parameter values to be properly set.
3. Publish the entire code necessary to quantify model performance. Specifically, the often lengthy post-processing and public resource integration steps should be included. Code versions should also be fixed and archived with Zenodo.

We additionally recommend researchers take advantage of modern package and environment management tools, whether that be a published Docker image with the exact software required built in or a tool like conda along with an environment file specifying the versions of all software used that can reproduce the environment (advice that we follow ourselves, see methods). By providing these easy-to-use tools, authors can ensure other researchers can set up their machines to reproduce and explore work with little or no effort. Secondly, all results should be produced in a programmatic manner to enable re-performing analysis and generating all figures by running a script. In addition to acting as an infallible protocol for the analysis, these scripts reduce the barrier for re-running analysis, which prevents mismatch between published parameters and the parameters actually used as it is less likely to leave behind stale results or make a mistake when updating compared to implicitly running commands.

These guidelines entail activities that are relatively easy for original authors to carry out, yet will substantially improve the velocity of the field, by decreasing wasted effort that goes into simply re-establishing baseline model performance, readily exposing flaws, and clarifying the relevance of sequence-to-expression models to disease.

## 4 Methods

### 4.1 Code

Code for Basenji and the released environment, `prespecified.yml,` were downloaded from https://github.com/calico/basenji on January 30, 2025. Note that the Basenji codebase as released by the authors is “something in between personal research code and accessible software for wide use,” [28]. It is not versioned to match the published results and contains many extra files that are not used. For readability, in our reproduction, we have reduced the modules to only those that are actually used. Code for running inference with Enformer was written by referencing released code available at https://github.com/google-deepmind/deepmind-research/tree/master/enformer.

We provide code to reproduce all our results, along with environmental files to make reproducing the analysis straightforward, and to avoid compatibility issues. We also supply more modular code to rerun specific analyses, as the full analysis is computationally intensive. To avoid conflicts with software already installed on other researcher’s systems, we supply a `Dockerfile`that can be used to build a `Docker image` containing each `conda` environment. The Dockerfile itself acts as a reference for setting up the `conda` environments. Using `Docker` allows a user to run code with minimal setup and without altering software already on their computer. For additional detail on these environments, we provide a `prespecified.yml` for each environment used in this paper. The `prespecified.yml` files provide information about software dependencies in a format the `conda` package manager can use to recreate each environment. Specifically, these files detail independent environments for running Basenji, Enformer, the GSEA, and for the main analysis.

We also supply all code outside of `Docker`, in an organized and modular fashion. To further simplify reproducing the results of this publication, we have written a `main.sh bash` script, which will run each piece of the analysis in order using the correct `conda` environment. Like the `Dockerfile,` even if the `main.sh` file is not used directly, it acts as a declarative list of commands run to complete the full analysis. Code used in this publication to retrain Basenji, run inference with Basenji and Enformer, compute performance and generate figures has been made available at https://github.com/adazhang1/genomics-replicability. Due to GitHub file size limits, model weights of the released models and retrained models are available at https://huggingface.co/adazhang1/genomics-replicability/tree/main/models and supporting datasets are available at https://huggingface.co/datasets/adazhang1/genomics-replicability/tree/main/supporting_data.

Detailed documentation on the code, supporting data, and how to train/test the models are available on both GitHub and Hugging Face to facilitate reproduction of the results in this publication. This code is versioned and archived at Zenodo: https://doi.org/10.5281/zenodo.20950355.

### 4.2 Calculating gene expression from CAGE

Gene annotations were obtained from GENCODE. The exact GENCODE file used is available in our `supporting_data` directory on the Hugging Face repository. It was processed using the `process_gtf_file` function and then cross-referenced with Basenji’s test sequences using the `locate_TSS_in_Basenji_tracks` function, both of which are implemented in our helper_functions.py file available on GitHub. In Basenji’s test data set, there were 1955 protein-coding genes. Ground truth and predicted gene expression values were obtained by first summing the cap analysis of gene expression (CAGE) read counts within the 128 bp bin containing each unique transcriptional start site (TSS) location of each protein-coding gene (and the two neighboring bins, i.e., 3 bins total), then computing a log_10_(1 + *x*) transformation on the summed read counts, and finally normalizing each CAGE track to have 0 mean and unit variance across genes via the `extract_gene_expression`. and normalize functions, also implemented in our `helper_functions.py` file available on GitHub.

### 4.3 Details of synthetic data sets modeled on properties of Enformer results

#### 1. Unimodal distribution

For a given reported correlation, one might initially imagine a unimodal Gaussian distribution with that correlation. To generate this distribution, we calculate the mean and variance of the ground truth and predicted gene expression for each CAGE track. The covariance between the ground truth and predicted values across the whole track were used in the off-diagonal entries of the covariance matrix. We then sampled *N* points from multivariate Gaussian *N* (***µ*, Σ**) where

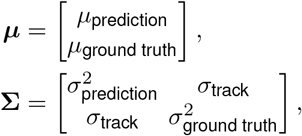

and *N* was the number of genes in each CAGE track

#### 2. Bimodal distribution where each mode uses the overall track correlation

Understanding that the distribution of gene expression might be bimodal, with one mode representing “Off” genes and one mode representing “On” genes, one might assume the correlation within the “On” and “Off” gene groups should be similar to the overall correlation. Using a threshold of -0.5 to separate the “On” and “Off” gene groups (as used in Fig. 1), for each CAGE track, we sampled *N*_ON_ points from the multivariate Gaussian *N* _ON_(***µ***_ON_, **Σ**_ON_) which was generated similarly to the unimodal distribution above, except that means and variances were computed for the “On” genes only, and *N*_ON_ was the number of genes in the “On” group. Note that the overall track covariance (as opposed to the covariance of the “On” genes) was used in the **Σ**_ON_ covariance matrix. I.e.,

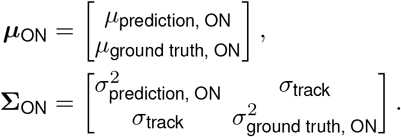

We similarly sampled *N*_OFF_ points from _OFF_(***µ***_OFF_, **Σ**_OFF_).

#### 3. Bimodal distribution where each mode has zero correlation

The correlation of each mode was surprisingly low compared to the overall correlation shown in Fig. 1. We therefore generated another bimodal distribution, similar to the one described above, but where each mode had zero covariance. I.e.,

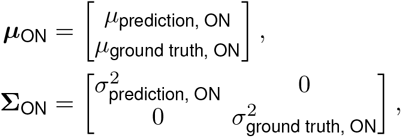

and similarly for the “Off” genes. This distribution represents the scenario where a model is only able to predict “On” or “Off”

Code for generating these synthetic datasets is implemented in the `generate_synthetic_data` function within our `helper_functions.py` file available on GitHub.

### 4.4 Gene popularity statistics

To determine the portion of scientific literature related to a given gene, we counted gene mentions in the publication network using the PubTator3 [70] dataset. PubTator3 uses a Name Entity Recognition (NER) model to automate detection of gene mentions across the PubMed database. We only used abstracts due to only a subset of all publications containing full text. Gene prevalence was defined as the number of publications in which a gene is mentioned. We stripped the Ensembl version numbers before mapping genes to Entrez identifiers used by PubTator to prevent mentions of a single gene from being split.

## Acknowledgments

This study was funded in part by the National Institutes of Health, R01AG061798.

