## Supplemental tables and figures for "Disease relevance and replicability of deep learning gene expression prediction"

| | Correlation | | "On"/"Off" Distance |
|---|---|---|---|
| | Basenji | Enformer | |
| Including cancer cells | 0.000433 | 0.000777 | 0.000002 |
| Excluding cancer cells | 0.000255 | 0.021435 | 0.000954 |

**Table S1:** *p*-values from ANOVA tests of tissue effect on model performance correlation and on GMM-computed distance between "On" and "Off" groups. Note that distance is computed only on ground truth expression, which makes it independent of model performance.

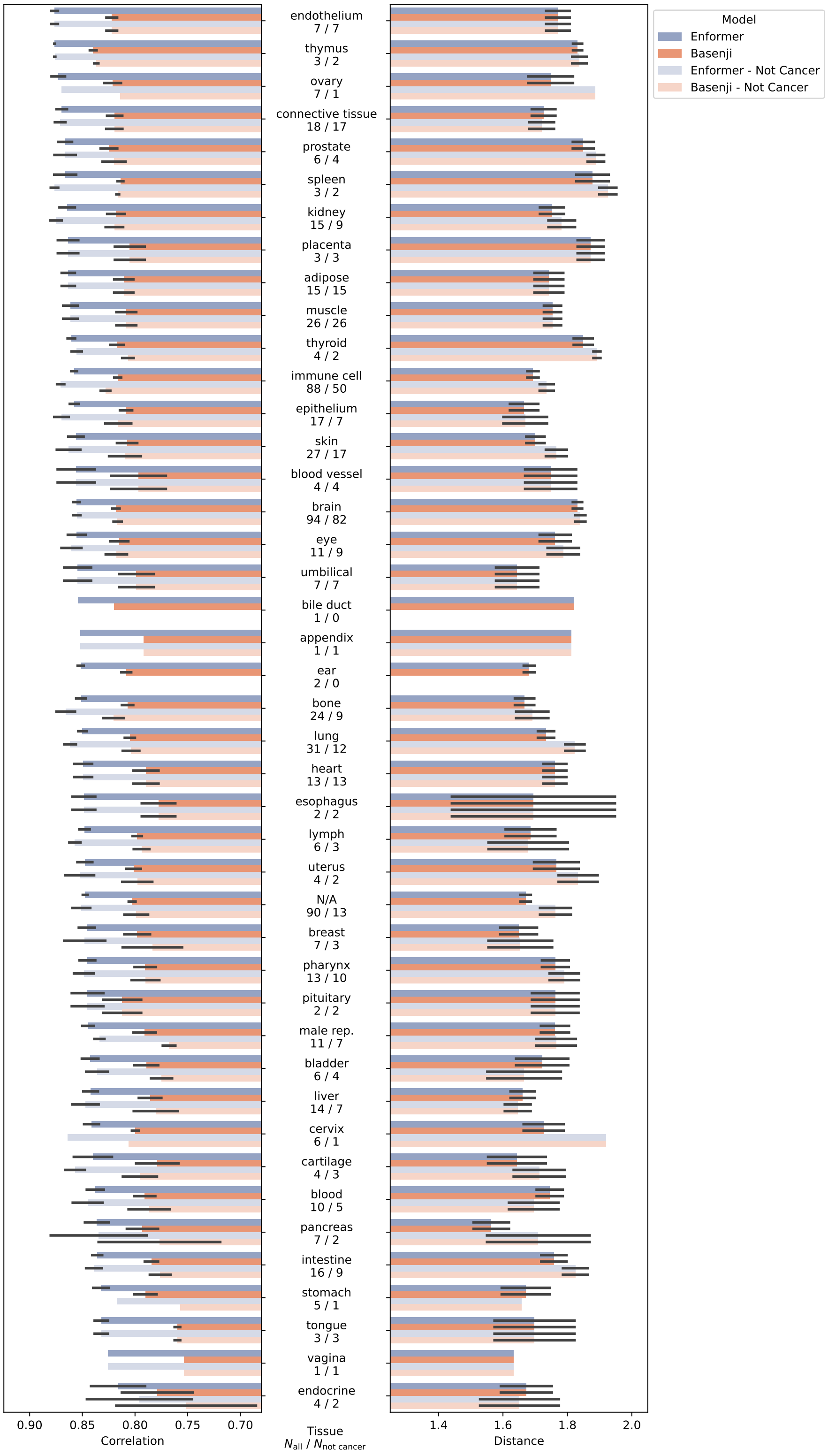


**Figure S1:** Mean correlation and GMM distance across CAGE tracks grouped by tissue type. Darker-colored bars are averages across all samples. Lighter-colored bars are averages excluding cancer lines. Number of CAGE tracks within a tissue type are also reported in the y-axis labels. Tissues are sorted by descending Enformer mean correlation when including cancer cells.

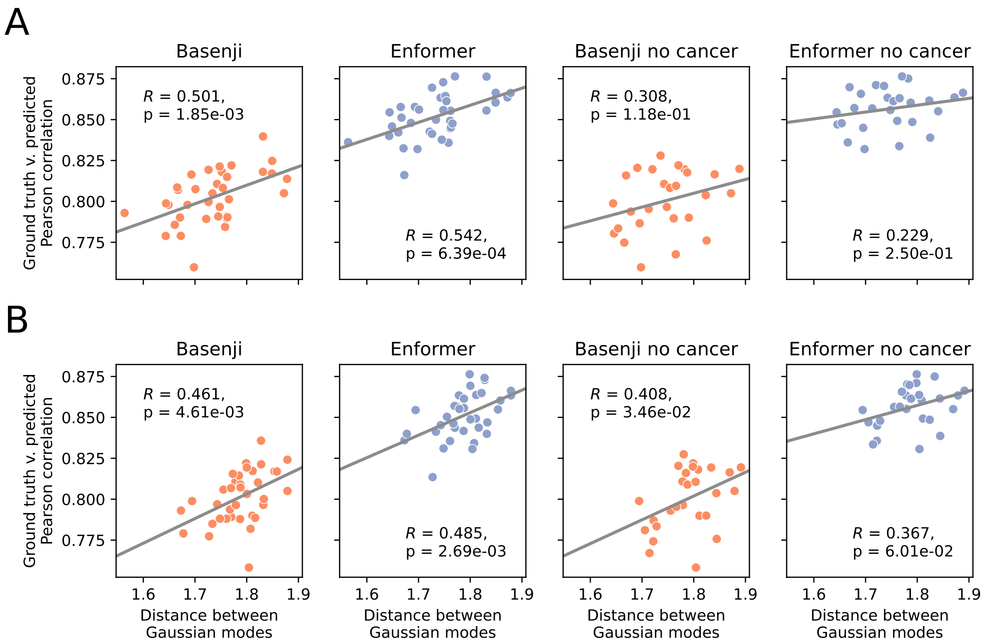


**Figure S2:** Tissue-based GMM distance versus correlation. Two methods for computing GMM distance and correlation were used. A) GMM distances and Pearson correlation for all CAGE tracks from a given tissue were averaged, then plotted. B) Ground truth values of all CAGE tracks for a given tissue were combined and a new GMM was fit to that distribution, and a new Pearson correlation was computed between the combined ground truth and predicted values.

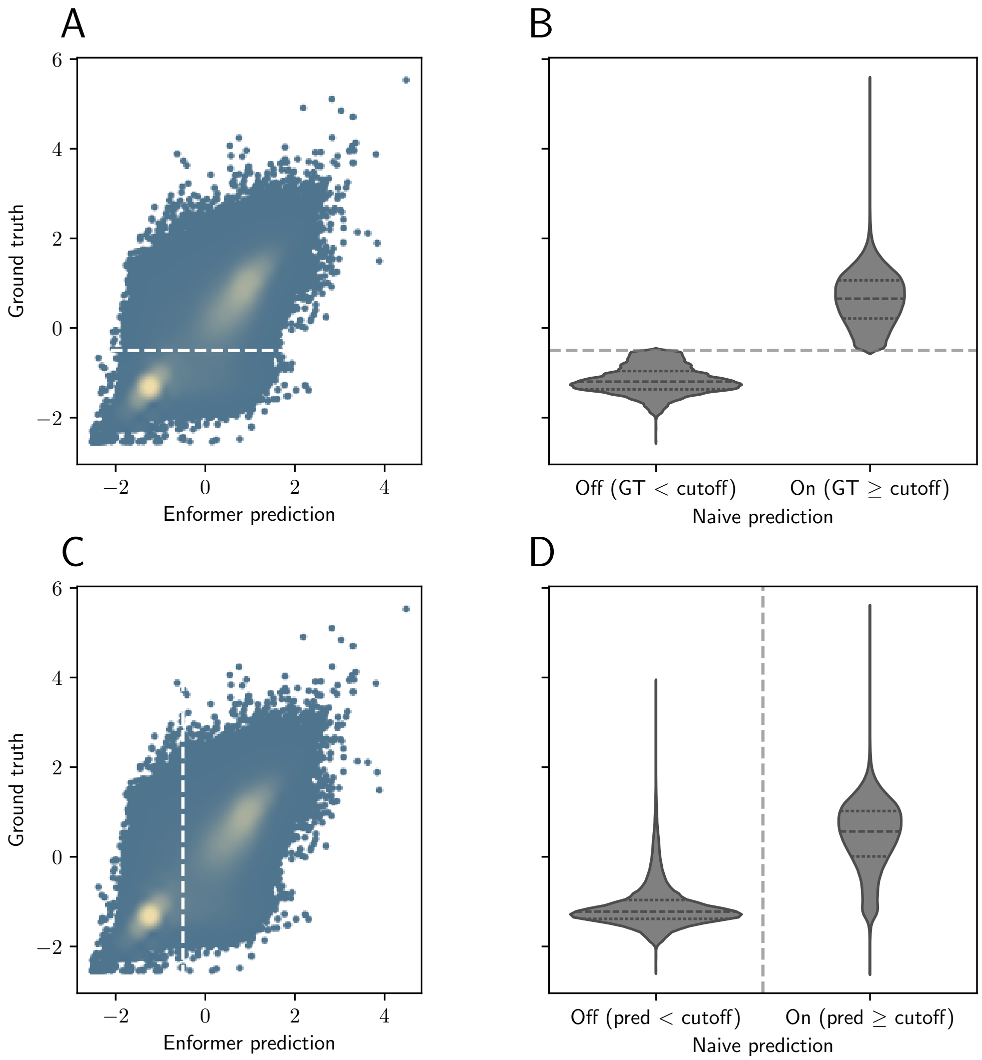


**Figure S3:** Generating the two "Naive" prediction models from Enformer. A) Generating a naive model which always returns a correct on or off results based on the ground truth. B) Violin plot of input values for the "On" or "Off" results for the ground truth model. C) naive model which always returns a correct "On" or "Off" results based on predicted values. D) Violin plot of input values for the "On" or "Off" results for the prediction-based naive model.

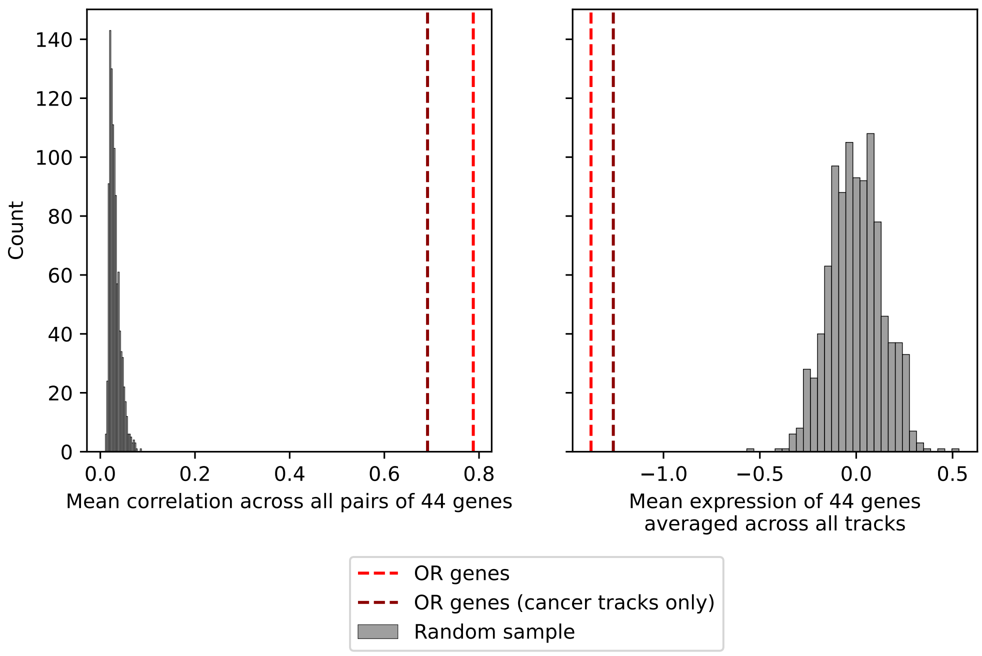


**Figure S4:** Comparing OR gene mean correlation between pairs and mean expression versus a random sample of the same number of genes.
